# Intestinal inflammation reversibly alters the microbiota to drive susceptibility to *Clostridioides difficile* colonization in a mouse model of colitis

**DOI:** 10.1101/2022.04.07.487579

**Authors:** Madeline R. Barron, Kelly L. Sovacool, Lisa Abernathy-Close, Kimberly C. Vendrov, Alexandra K. Standke, Ingrid L. Bergin, Patrick D. Schloss, Vincent B. Young

## Abstract

Susceptibility to *Clostridioides difficile* infection (CDI) typically follows the administration of antibiotics. Patients with inflammatory bowel disease (IBD) have increased incidence of CDI, even in the absence of antibiotic treatment. However, the mechanisms underlying this susceptibility are not well understood. To explore the intersection between CDI and IBD, we recently described a mouse model where colitis triggered by the murine gut bacterium, *Helicobacter hepaticus,* in IL-10^-/-^ mice led to susceptibility to *C. difficile* colonization without antibiotic administration. The current work disentangles the relative contributions of inflammation and gut microbiota in colonization resistance to *C. difficile* in this model. We show that inflammation drives changes in microbiota composition, which leads to CDI susceptibility. Decreasing inflammation with an anti-p40 monoclonal antibody promotes a shift of the microbiota back toward a colonization-resistant state. Transferring microbiota from susceptible and resistant mice to germ-free animals transfers the susceptibility phenotype, supporting the primacy of the microbiota in colonization resistance. These findings shine light on the complex interactions between the host, microbiota, and *C. difficile* in the context of intestinal inflammation, and may form a basis for the development of strategies to prevent or treat CDI in IBD patients.

**Importance:** Patients with inflammatory bowel disease (IBD) have an increased risk of developing *C. difficile* infection (CDI), even in the absence of antibiotic treatment. Yet, mechanisms regulating *C. difficile* colonization in IBD patients remain unclear. Here, we use an antibiotic-independent mouse model to demonstrate that intestinal inflammation alters microbiota composition to permit *C. difficile* colonization in mice with colitis. Notably, treating inflammation with an anti-p40 monoclonal antibody, a clinically relevant IBD therapeutic, restores microbiota-mediated colonization resistance to the pathogen. Through microbiota transfer experiments in germ-free mice, we confirm that the microbiota shaped in the setting of IBD is the primary driver of susceptibility to *C. diffiicile* colonization. Collectively, our findings provide insight into CDI pathogenesis in the context of intestinal inflammation, which may inform methods to manage infection in IBD patients. More broadly, this work advances our understanding of mechanisms by which the host-microbiota interface modulates colonization resistance to *C. difficile*.

## Introduction

The mammalian gastrointestinal tract is inhabited by a diverse community of microbes that contributes to colonization resistance against pathogenic organisms, including the toxin-producing bacterium, *Clostridioides difficile* (1, 2). Disruption of the microbiota, typically in the setting of treatment with antibiotics, allows *C. difficile* to establish within the gut (3–7). *C. difficile* colonization can lead to clinical syndromes ranging from mild diarrhea to severe colitis (8). Host immune responses also support colonization resistance to the pathogen, in part through modulating the structure and function of the gut microbial community (9–11).

The interplay between the host and microbiota also underlies the pathogenesis of the inflammatory bowel diseases (IBD) Crohn’s disease and ulcerative colitis. IBD results from dysregulated interactions between host immune responses and the microbiota in genetically susceptible individuals, leading to chronic, progressive inflammation of the gut (12). IBD patients harbor a distinct microbiota compared to healthy individuals (13). These changes in microbiota structure and associated inflammation in IBD are accompanied by an altered intestinal metabolome, characterized by decreases in short chain fatty acid concentrations and altered bile acid profiles (14–16). Notably, patients with IBD have an increased risk of developing *C. difficile*infection (CDI) (17–21), even in the absence of antibiotic treatment (22).

However, despite the known associations between inflammation and microbiota alterations in IBD, the mechanisms permitting colonization by *C. difficile* in IBD patients remain unclear.

To explore the intersection between CDI susceptibility and IBD, we developed a mouse model in which *C. difficile* colonization occurs in the absence of antibiotic pretreatment (10). This is in marked contrast to most models of CDI, where antibiotic administration is required to make animals susceptible to infection (4, 23, 24). In our novel system, IL-10^-/-^ mice are colonized by *Helicobacter hepaticus,* a murine gut bacterium that triggers colitis in genetically predisposed hosts (25, 26). The combination of intestinal inflammation, coupled with changes in microbiota composition, is sufficient to render colitic mice susceptible to colonization by *C. difficile* (10). However, the relative contributions of inflammation and microbiota alterations underlying susceptibility to *C. difficile* in this model remain to be explored.

In the current study, we sought to separate the differential role of host responses and changes to the microbiota that lead to loss of resistance to *C. difficile* colonization in mice with colitis. To do this, we follow up on observations that the pro-inflammatory cytokine IL-23 drives *H. hepaticus*-triggered colitis in mice with defective regulatory immune signaling (27, 28). We show that inflammation changes the composition of the gut microbiota to permit *C. difficile* colonization. Subsequently, treating inflammation with a monoclonal antibody that targets the p40 subunit of IL-23 restores resistance to *C. difficile.*Transferring the microbiota from colonization resistant or inflammation-induced susceptible IL-10^-/-^ mice to germ-free wildtype mice is sufficient to transfer the susceptibility phenotype, underscoring the central role of the microbiota in colonization resistance to *C. difficile*. Together, our results provide insights into mechanisms by which the host-microbiota interface mediates susceptibility to *C. difficile* colonization in the context of intestinal inflammation.

## Materials and Methods

### Bacterial culture

*Helicobacter hepaticus* strain 3B1 (ATCC 51448) was streaked onto tryptic soy agar (TSA; Hardy Diagnostics) blood plates containing 5% sheep’s blood and grown for 3-4 days in a vinyl hypoxic chamber (2% O_2_, 5% CO_2;_ Coy Industries) maintained at 37°C. For murine inoculations, *H. hepaticus* from actively growing plates was streaked onto fresh blood agar and grown an additional 3-4 days. Each plate was washed with sterile tryptic soy broth (TSB; Spectrum Chemical) and bacterial suspensions were combined and centrifuged at 4,000xg for 10 mins. The resulting pellet was re- suspended in fresh TSB for oral gavage of mice.

To prepare *C. difficile* spore stocks, *C. difficile* VPI10463 was grown overnight at 37°C in 5 mL of Columbia Broth in a vinyl anaerobic chamber (Coy Industries). Three overnight cultures were prepared. The following day, the cultures were each added to 40 mL of Clospore media (29) and incubated at 37°C for 14 days. After this time, the tubes were centrifuged at 4°C and washed 2x in ice-cold sterile water. After another wash in cold 0.05% Tween 20, the pellets were combined and washed again. The pellet was suspended in 1 mL of cold 50% Histodenz™ gradient (Sigma Aldrich) diluted in distilled, nuclease-free water and carefully pipetted into a conical tube containing an additional 19mL of Histodenz™. After centrifugation for 20 mins, the spores were washed with 0.05% Tween 20 once, then ice-cold water 3 more times. The final pellet was suspended in water and transferred to a microcentrifuge tube for storage at 4°C. Spore purity was verified via phase-contrast microscopy. Spore inocula for mouse infections were prepared as previously described (24). The inocula were quantified via serial plating on pre-reduced cycloserine-cefoxitin-fructose agar containing 0.1% taurocholate (TCCFA), both at the time of preparation (3 days before challenging mice) and after infection. TCCFA was prepared as previously described (30).

### Mice

All animal experiments were completed with approval from the University of Michigan Committee on Use and Care of Animals. Male and female C57BL/6 IL-10- deficient mice, aged 6-11 weeks at the start of experiments, were used. Co-housed littermates of same sex were randomly assigned to experimental groups. All mice were from a breeding colony obtained from Jackson Laboratories over 20 years ago. Animals were housed in specific-pathogen-free (SPF), *Helicobacter*-free conditions at 22°C under a 12h light/dark cycle, and animal husbandry was performed in an AAALAC-accredited facility. For microbiota transfer experiments, male and female germ-free C57BL/6Ncr mice aged 7-11 weeks were obtained from a colony established and maintained by the University of Michigan Germ-Free Mouse Facility. Co-housed littermates of same sex were randomly assigned to experimental groups and mice received sterile food, water, and bedding throughout the experiments.

### Helicobacter hepaticus colitis

To trigger colitis in IL-10^-/-^ mice, animals were inoculated with ∼10^8^ colony- forming units (CFU) of *H. hepaticus* in 100 uL volume TSB via oral gavage (31). Control mice received sterile TSB. *H. hepaticus* colonization was confirmed by PCR with primers specific for the *H. hepaticus* 16S rRNA gene (32) (Table S1) using DNA isolated from feces 5-7 days post-inoculation. For extractions, feces were homogenized in 500 uL sterile PBS (Gibco) and the homogenate centrifuged at 500xg for 10-15 secs. DNA was extracted from the resulting supernatant using DNeasy UltraClean Microbial Kit (Qiagen), following manufacturer’s instructions.

### Monoclonal antibody treatments

Anti-p40 (Bio X Cell Cat# BE0051, RRID: AB_1107698) and mouse IgG2A isotype control (Bio X Cell Cat# BE0089, RRID: AB_1107769) monoclonal antibodies were diluted in *InVivo*Pure™ pH 7.0 Dilution Buffer (Bio X Cell). Mice were injected intraperitoneally with 1mg of antibody suspended in 200 uL buffer every 3-4 days for 3 weeks, beginning 14 days post-*H. hepaticus* inoculation. The frequency of injections was determined based on preliminary studies. Mice without colitis received 200 uL of dilution buffer at each timepoint. Mice challenged with *C. difficile* continued to receive antibodies after infection until the end of experiment.

### *C. difficile* infections and quantification from intestinal content

Animals were administered ∼ 3x10^4^ *C. difficile* strain VPI10463 spores in 40-50uL water or mock-challenged with water by oral gavage. Mice were monitored for signs of clinical disease. Disease scores were averaged based on scoring of the following features for signs of disease: weight loss, activity, posture, coat, diarrhea, and eyes/nose. A 4-point scale was assigned to score each feature, and the sum of these scores determined the clinical disease severity score (33).

To measure *C. difficile* colonization, fecal pellets were collected in pre-weighed sterile tubes. After collection, tubes were re-weighed to determine fecal weight, and passed into a vinyl anaerobic chamber (Coy Industries). Feces were diluted 1:10 (w/v) in pre-reduced sterile PBS (Gibco). The fecal homogenate was serially diluted in PBS and 100 uL was plated on pre-reduced TCCFA; the same procedure was followed for plating cecal contents at necropsy. TCCFA plates with fecal or cecal samples were incubated at 37°C for at least 18 hours prior to colony enumeration.

### Clinical disease severity and histopathology

At the end of experiments, mice were euthanized via CO_2_ asphyxiation and colon and cecal tissues were collected and fixed in 10% formalin and stored in 70% ethanol until processing. Tissue was embedded in paraffin and sectioned to generate hematoxylin and eosin-stained slides. To assess colitis severity resulting from *H. hepaticus* colonization, slides were scored for lymphocytic inflammation by a board-certified veterinary histopathologist (I.L.B.) blinded to the experimental groups using the following a 4-point scoring system, as previously described (34). Histopathologic damage associated with *C. difficile* infection was scored using epithelial destruction, immune cell infiltration, and edema on 4-point scale and the sum of all categories was used to determine histological score (24, 35).

### Microbiota transfer experiments

Inocula for microbiota transfers (MT) were prepared as follows. Fecal pellets were collected in pre-weighed tubes from donor mice one day prior to challenge with *C. difficile* and frozen at -80°C. Donors included colitic IL-10^-/-^ mice treated with control mAb or anti-p40 mAb that went on to become susceptible or remain resistant to *C. difficile* colonization. On the day of MT, donor fecal pellets were thawed, re-weighed, and passed into an anaerobic chamber. Feces were diluted 1:20 (w/v) in pre-reduced PBS (Gibco) and vortexed for 10 mins at maximum speed. Samples were centrifuged at 800rpm for 30 secs. 50-70 uL of the supernatant was administered to germ-free animals via oral gavage. After inoculation, the gavage material was plated on blood agar plates and incubated overnight in an anaerobic chamber (37°C) to confirm the presence of viable bacteria. After 7 days, mice were challenged with *C. difficile* and colonization monitored as described above.

### Fecal lipocalin-2 quantification

Fresh fecal pellets were collected from mice at baseline, two weeks post-*H. hepaticus* inoculation, and after 3 weeks of monoclonal antibody treatment in pre- weighed microcentrifuge tubes. Tubes were re-weighed to determine fecal weight and the pellet suspended in PBS (Gibco) with 0.1% Tween 20. Suspensions were vortexed at maximum speed for 20 mins to homogenize the sample and centrifuged at 4°C for 10 mins. The supernatant was collected and used to quantify fecal lipocalin-2 levels using the mouse lipocalin-2/NGAL DuoSet ELISA kit (R&D Systems) in a microplate reader according to manufacturer’s instructions.

### Tissue RNA extraction and RT-PCR

Colon tissue snips collected at time of necropsy were placed in RNA*later*™ solution (Invitrogen) and stored overnight at 4°C, then transferred to -80°C until processing. Tissues were weighed, homogenized, and total RNA was extracted from samples using the AllPrep DNA/RNA Mini Kit (Qiagen) according to manufacturer’s instructions. After extraction, RNA was diluted to 50 ng/uL for RTPCR reactions.

For one-step RT-PCR analyses, 25 uL reaction mixtures were prepared using the TaqMan® RNA-to-Ct™ 1-Step Kit (Thermo Fisher) per manufacturer instructions, with 150 ng of RNA used per reaction. All reactions were run in technical duplicates with appropriate controls. RT-PCR was performed on a LightCycler96 qPCR machine (Roche) with initial incubations at 48°C for 15 mins and 95°C for 10 mins, followed by 40 cycles of 95°C for 15s and 60°C for 1 min. Relative expression of targets was determined via the 2 ^ΔΔ*CT*^ method, using β actin as the control gene. Primers used for 1-step RT-PCR are depicted in Table S1. All probes were modified with FAM and TAMRA at the 5’ and 3’ ends, respectively.

### Targeted metabolomics

Quantification of cecal short chain fatty acids (SCFA) and bile acids was completed by the University of Michigan Medical School Metabolomics Core. Cecal contents were collected in pre-weighed sterile tubes, immediately frozen in liquid nitrogen, and stored at -80°C until submission. Prior to submission, tubes were re- weighed to determine weight of cecal contents, and approximately 50 mg submitted for use in subsequent assays.

*SCFAs:* Water and acetonitrile containing internal standards (500 µM D4-acetic acid, 250 µM D7-butyric acid, and 6.25 µM D11-hexanoic acid) were added to each sample. Samples were homogenized (Branson) and centrifuged at 15,000xg at 4°C for 10 mins. Supernatant was transferred to a 1.8 mL glass autosampler vial and 200mM 3- nitrophenylhydrazine (3-NPH) in 1:1 acetonitrile: water and 120 mM of 1-Ethyl-3-(3- dimethylaminopropyl) carbodiimide in 1:1 acetonitrile water with 6% pyridine were added. Samples were vortexed and placed in a warming oven at 40 °C for 30 mins.

Once derivatization was complete, samples were cooled, and 90/10 water/acetonitrile was added. Standards were prepared identically to cecal samples, substituting volatile fatty acid mix (Sigma Aldrich) diluted to concentrations ranging from 3 µM to 3000 µM. Samples were analyzed via liquid chromatography mass spectrometry (LC-MS) using an Agilent 1290 LC coupled to an Agilent 6490 triple quadrupole MS and a Waters HSS T3, 2.1 mm x 100 mm, 1.8 µm particle size chromatographic column. Mobile phase A contained 0.1% formic acid in water while mobile phase B consisted of 0.1% formic acid in methanol. The gradient was as follows: linear ramp from 15% to 80% B from 0-12 min; step to 100% B from 12-12.1 min; hold 100%B from 12.1-16 min; step to 15% B from 16-16.1 min; hold 15% B from 16.1- 20min. The injection volume was 5 µL and the column temperature was 55 °C. MS parameters were as follows: gas temp 325°C, gas flow 10 L/min, nebulizer 40 psi, capillary voltage 4000V, scan type MRM, negative ion mode, delta EMV 600.

Quantitation was performed using MassHunter Quantitative Analysis software (Agilent v.B.07.00) by measuring the ratio of peak area of the 3-NPH derivatized SCFA species to its closest internal standard (by retention time). Linear standard curves were used to estimate SCFA concentrations in the extract, which were normalized to the measured mass of cecal contents.

*Bile acids:* To determine bile acid concentrations, extraction solvent containing chilled acetonitrile with 5% NH_4_OH and isotope-labeled internal standards were added to 20 mg of cecal samples. Samples were homogenized by probe sonication for 20 secs. The homogenized mixture was centrifuged, and an aliquot of supernatant transferred to an LC-MS autosampler and dried in a speedvac set to 45°C for approximately 45 mins.

Samples were reconstituted in 50/50 methanol/water. A series of calibration standards ranging from 0 to 1000 nM were prepared along with samples for metabolite quantification.

Bile acid LC-MS analyses were performed on an Agilent 1290 LC coupled with a 6490 Triple Quad mass spectrometer operated in MRM mode. Metabolites were separated on a HSS T3, 2.1 mm x 50 mm, 1.8m particle size column using water + 0.1% formic acid as mobile phase A, and acetonitrile + 0.1% Formic acid, as mobile phase B. The flow rate was 0.35 mL/min with the following gradient: linear from 5 to 25% B over 2 mins, linear from 25 to 35% B over 16 mins, linear from 35 to 75% B over 8 mins, followed by isocratic elution at 95% B for 8 mins. The system was returned to starting conditions (5% B) in 0.1 min and held there for 3 mins to allow for column re-equilibration before injecting another sample. Data were processed using MassHunter Quantitative analysis. Metabolites were quantitated by determining the ratio of the peak area for each compound to that of the closest-matching internal standard (by RT), and then calculating concentration using a 5-point linear calibration curve prepared using authentic standards. Calibration curve accuracy was determined to be better than 80% for each standard. Measured bile acid concentrations were normalized to dry sample weight after quantification.

### 16S rRNA-encoding gene amplicon sequencing and analysis

The University of Michigan Microbiome Core extracted total DNA from intestinal contents, including either whole fecal pellets or 200-300 uL of contents diluted 1:10 (w/v) in sterile PBS using the MagAttract PowerMicrobiome Kit (Qiagen), and prepped DNA libraries as previously described (36). Samples were randomized into each extraction plate. DNA was amplified using dual-index primers targeting the V4 region of the 16S rRNA gene, as described previously (37). Sequencing was conducted on the Illumina MiSeq platform using the MiSeq Reagent kit V3 for a total of 500 total cycles, with modifications found in the Schloss SOP (https://github.com/SchlossLab/MiSeq_WetLab_SOP).

To assess sequencing error, the V4 region of the ZymoBIOMICS Microbial Community DNA standard (Zymo Research) was also sequenced. Data were analyzed using mothur (v.1.44.2) software package (38). Briefly, following assembly, filtering, and trimming, contigs were aligned to the Silva v.128 16S rRNA database. Any sequences that failed to align, or were flagged as possible chimeras by UCHIME, were removed (39). Sequences were clustered into operational taxonomic units (OTUs) with Opticlust (40) using a 97% similarity cut-off and classified via a Bayesian classifier using the Silva rRNA database. LEfSe analysis was conducted in mothur using the “lefse” command with default settings. The limit of detection for relative abundance analyses was calculated as the smallest non-zero relative abundance value observed in the dataset divided by 10.

### Computational modeling and machine learning analyses

Dirichlet multinomial mixutures (DMM) modeling (41) was used to achieve an unbiased analysis of the association between the microbiota at the time *C. difficile* first contacts the gut environment, and downstream susceptibility or resistance to the pathogen in mice with IBD treated with control mAb or anti-p40. Analyses included samples from 3 independent experiments, two in which mice were harvested day 9 post-*C. difficile* challenge, and one where animals were sacrificed one day post-challenge. Animals were classified based on whether they were positive for *C. difficile* one day post-challenge to include samples from all experiments. Additionally, temporal colonization experiments revealed that positivity for *C. difficile* one day post-challenge was associated with a long-term colonization phenotype, suggesting that this was a strong readout of robust colonization. DMM was completed in mothur using the “get.communitytype” command with default settings.

Supervised machine learning was performed according to the best practices outlined by Topçuoğlu et al., 2020 and implemented in the mikropml R package v1.2.1 (42). Models were trained on relative abundance data from fecal samples collected from animals on the day of *C. difficile* challenge predict presence of the pathogen on day 1 post-challenge. The data were first pre-processed by centering and scaling abundance counts, collapsing perfectly correlated OTUs, and removing OTUs with zero variance.

For 100 random seeds, the data were randomly split into training and testing sets with 65% and 35% of the samples in each, respectively. Random forest models were trained on the training sets using 5-fold cross-validation to select the best hyper-parameter value (mtry: the number of OTUs included per tree), then the best models were evaluated on the held-out test sets by computing the AUROC and AUPRC. AUPRC is a useful metric for evaluating binary classifiers when there is an imbalance in the number of positive and negative events in a dataset (i.e., a larger fraction of animals negative for *C. difficile* day 1 post-challenge compared to positive) (43). An AUROC of 1 indicates the model perfectly distinguishes between sample groups, while an AUROC of 0.5 indicates the model does not predict better than random chance. For AUPRC, the baseline performance is calculated as the number of positive samples over the total number of samples, or, in this case, 0.34.

The most important OTUs contributing to model performance were determined by permutation feature importance tests (44). First, perfectly correlated OTUs were collapsed into a single representative. Then for each trained model, each OUT in the test dataset was randomly shuffled 100 times and the new permutation performance (AUROC) was measured. A given OUT was considered significantly important for a model at an alpha level of 0.05, where less than 5% of the permutation AUROC values were greater than the original test AUROC. The OTUs that decreased the AUROC the most when permuted were considered the most important for model performance.

### Statistical analyses

Statistical analyses were performed in Prism (GraphPad Software) or R. For comparing non-normally distributed data a Kruskal Wallis test followed by Dunn’s multiple comparisons test, or Mann Whitney U test, were performed. Data with normal distribution were analyzed using an ANOVA coupled with a Tukey’s post-hoc test. We employed a Chi-square test to determine the relationship between *C. difficile* colonization susceptibility and IBD treatment group. A Fisher’s exact test revealed correlations between microbiota enterotype, treatment group, and *C. difficile* positivity. Differences in Theta YC distances between MT recipients that were susceptible or resistant to *C. difficile* were analyzed using analysis of molecular variance (AMOVA). For all analyses, a P value of less than 0.05 was considered statistically significant.

Adobe Illustrator CC 2020 was used to arrange panels, modify color schemes as needed, and generate final figures.

### Data availability

The workflow used to perform the machine learning analysis is available at https://github.com/SchlossLab/Barron_IBD-CDI_2022. Data and code for remaining microbiota analyses can be found at https://github.com/barronmr/antip40_microbiota.

Raw 16S rRNA sequences have been deposited in the NCBI Sequence Repository Archive under the accession number PRJNA811422.

## Results

Treatment of inflammation in colitic mice by anti-p40 monoclonal antibody inhibits susceptibility to *C. difficile* colonization

In our previous work, we hypothesized that inflammation promotes susceptibility to *C. difficile* colonization in mice with colitis triggered by *H. hepaticus* (10). To further explore this hypothesis, we sought to determine whether treating established colitis would restore colonization resistance to *C. difficile* in this model.

C57BL/6 IL-10^-/-^ SPF mice were inoculated with *H. hepaticus* while non-IBD control animals received sterile tryptic soy broth via oral gavage (Fig. 1A). Intestinal inflammation was monitored by measuring fecal levels of the inflammatory marker, lipocalin-2 (Fig. 1A-B). Animals colonized with *H. hepaticus* developed colitis, as indicated by a significant increase in fecal lipocalin-2 concentrations relative to control mice two weeks after inoculation with *H. hepaticus* (Fig. 1B; day -21). At this time, anti- p40 monoclonal antibody (mAb), an isotype control mAb, or mAb vehicle was administered to animals by intraperitoneal injection every 3-4 days for 3 weeks (Fig. 1A). After 3 weeks of mAb administration, there was a reduction in lipocalin-2 concentrations in mice treated with anti-p40 mAb while levels remained elevated in mice that received the isotype control mAb (Fig. 1B). Histopathologic analysis of colonic tissue collected from mice after 3 weeks of antibody treatment (Fig. S1A) supported a significant decrease in inflammation in mice treated with anti-p40 mAb (Fig. S1B).

**Figure 1.**
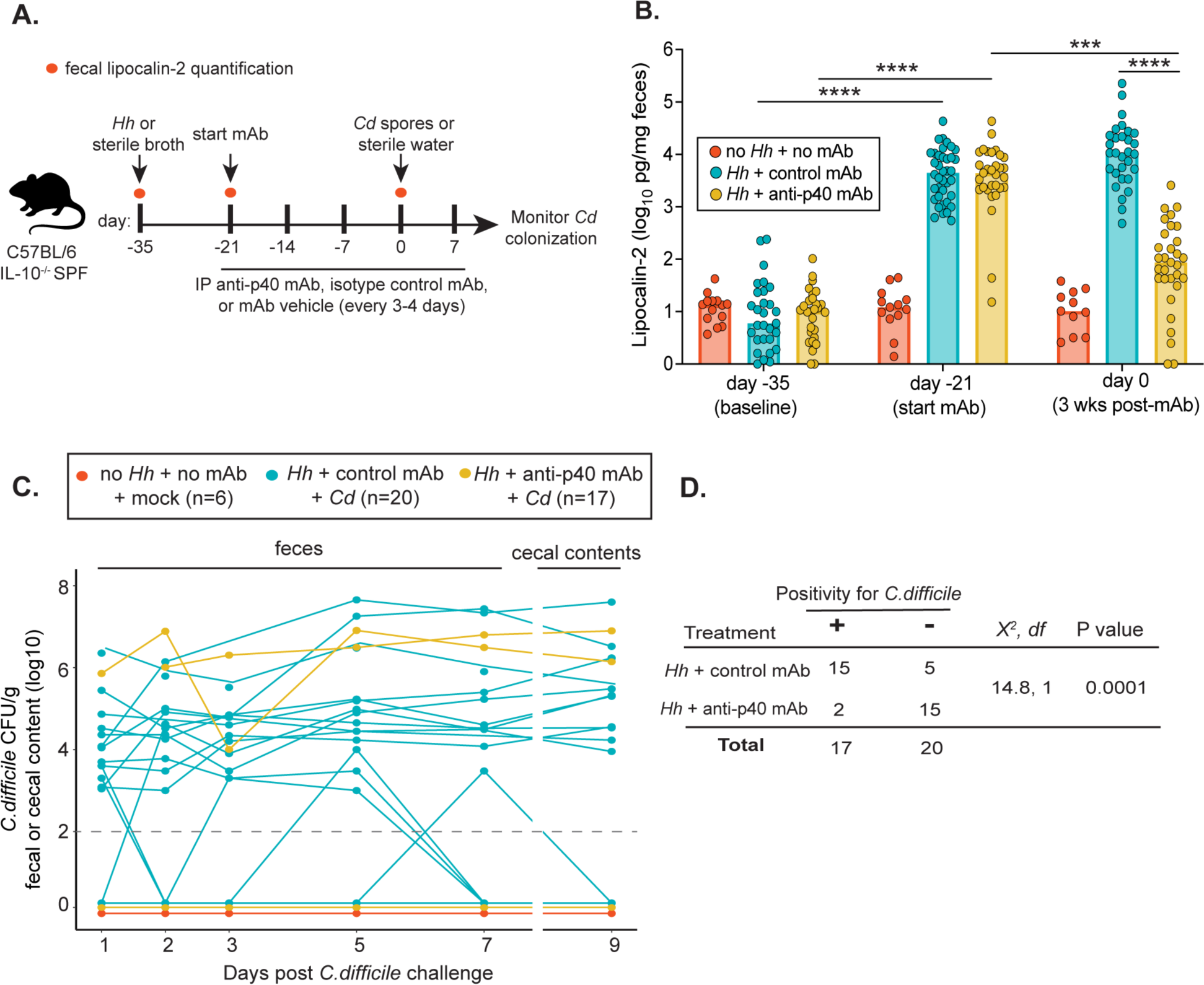
Treatment of inflammation in colitic mice by anti-p40 monoclonal antibody inhibits susceptibility to *C. difficile* colonization. (A) Experimental design. Mice were inoculated with *H. hepaticus* or sterile broth (non-IBD controls) via oral gavage. On day -21, animals were administered anti-p40 mAb or isotype control mAb via intraperitoneal (IP) injection every 3-4 days for 3 weeks. Non-IBD control animals received mAb vehicle. Intestinal inflammation before and after colitis development, and after 3 weeks of mAb treatment, was monitored by measuring fecal levels of lipocalin-2. Mice were then challenged with spores from *C. difficile* strain VPI10463 (day 0) and *C. difficile* colonization monitored over time. Control animals were mock challenged with sterile water. *Hh = H. hepaticus, Cd = C. difficile.* (B) Lipocalin-2 levels were measured via ELISA using feces collected from mice at baseline (day -35), after the development of IBD (day -21, first day of mAb treatment) and after 3 weeks of treatment with mAb or vehicle (day 0 of *C. difficile* challenge). Data are from 3 independent experiments. A Kruskal Wallis test followed by the Dunn’s multiple comparison’s test was performed (***P<0.001, ****P<0.0001). (C) Shedding of *C. difficile* in feces collected from animals over time, and in cecal contents collected at harvest (day 9). Positivity was determined as having viable *C. difficile* in intestinal contents after plating on TCCFA plates. The dashed line refers to the limit of *C. difficile* colonization detection (10^2^ CFU/g). Each line in the graph refers to a single mouse; lines below the limit of detection include multiple mice within each group. Data are from 2 independent experiments. (**D)** Chi-square analysis of mice with IBD depicted in (**C**) treated with isotype control mAb or anti-p40 mAb that were positive or negative for *C. difficile* at any point throughout the experiments. See also Figure S1.

Examination of these tissues revealed reduced hyperplasia and restoration of goblet cell mass in anti-p40 mAb-treated animals relative to those treated with control mAb (Fig. S1C). Moreover, colonic expression of genes encoding the cytokines IL-17A, IFN-γ, TNF-α, and IL-22 was also significantly lower in mice treated with anti-p40 relative to control mAb-treated animals (Fig. S1D-G). Thus, anti-p40 mAb effectively mitigates colitis in IL-10^-/-^ animals from our breeding colony, as previously reported (27, 28). To test the hypothesis that treating inflammation restores resistance to *C. difficile* colonization, mice were challenged with either ∼3x10^4^ spores of *C. difficile* strain VPI10463 or sterile water via oral gavage after 3 weeks of mAb treatment (Fig. 1A; day 0). Animals were monitored for *C. difficile* colonization and disease and continued to receive mAb injections throughout the rest of the experiment (Fig. 1A).

One day after spore challenge, 13/20 (65%) of mice that had received the isotype control mAb shed *C. difficile* in their feces (Fig. 1C). By 9 days after spore challenge, 15/20 (75%) of these mice had shed *C. difficile* at some point throughout the experiment compared to 2/17 (11.8%) mice treated with anti-p40 mAb (Fig. 1C-D). Thus, colitic mice treated with the control mAb were significantly more susceptible to *C. difficile* colonization than those treated with anti-p40 mAb (P=0.0001 by chi-square) (Fig. 2D). Following challenge with *C. difficile*, signs of clinical disease were mild in mice from both groups. Clinical disease scores did not differ between animals with IBD that had shed *C. difficile* during the experiment compared to animals that never shed *C. difficile*, regardless of which mAb they received (Fig. S2A). Similarly, histopathological analysis of cecal and colon tissue did not reveal a significant difference in the degree of edema, inflammatory infiltrate, and epithelial damage in *C. difficile*-susceptible and resistant mice within each treatment group (Fig. S2B), despite the reduction of inflammation in animals that received anti-p40 mAb. Collectively, these results confirm that inflammation creates a permissive landscape for *C. difficile* colonization and, when inflammation is decreased via anti-p40 mAb treatment, susceptibility to *C. difficile* is lost.

**Figure 2.**
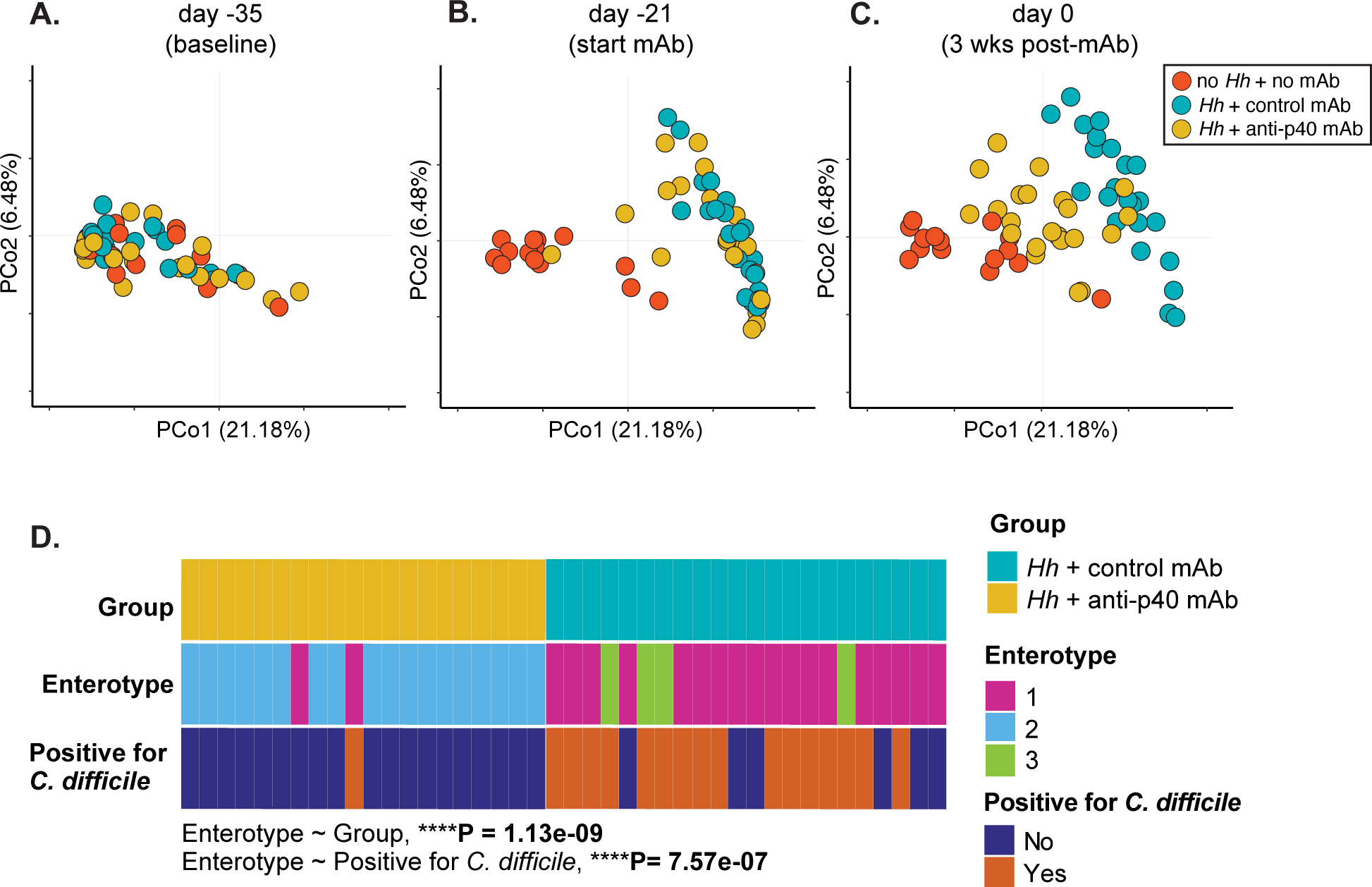
IBD-induced changes in microbiota structure are associated with susceptibility to *C. difficile* colonization. (**A-C**) Principal component analysis plot of Theta YC distances of bacterial communities in feces collected from mice at (**A**) baseline (day -35), (**B**) 21 days prior to challenge with *C. difficile* (day -21; start of mAb treatment) and (**C**) day 0 of *C. difficile* challenge (after 3 weeks of mAb treatment). Data are from 3 independent experiments. *Hh= H. hepaticus; Cd= C. difficile.* **(D**) Dirichlet multinomial mixtures modeling of 16S rRNA sequences from feces collected from mice with IBD treated with control mAb or anti-p40 mAb on day 0 of *C. difficile* challenge. Samples are from 3 independent experiments. A Fisher’s exact test was used to determine associations between enterotype and treatment group, and whether animals became positive for *C. difficile* on day 1 post-challenge. See also Figure S2.

### IBD-induced changes in microbiota structure are associated with susceptibility to ***C. difficile* colonization**

Given alterations in gut microbiota composition underlie the pathogenesis of both IBD and CDI (12, 45), we used 16S rRNA-encoding gene amplicon sequence analysis to examine the fecal microbiota of mice prior to *H. hepaticus* colonization (baseline), 21 days before challenge with *C. difficile* (when inflammation has developed and immediately prior to initiation of mAb treatment), and on the day of *C. difficile* challenge (after 3 weeks of mAb treatment). The microbiota of groups that were treated with mAb were compared to animals that were not colonized with *H. hepaticus* or treated with mAb. At baseline, all mice had a similar gut microbiota structure (Fig. 2A). The development of colitis was accompanied by alterations in microbial community composition (Fig. 2B). By the day of *C. difficile* challenge, and following 3 weeks of mAb treatment, the microbiota of anti-p40 mAb-treated animals had shifted toward baseline (Fig. 2C). A similar trend was observed in the cecal contents of animals harvested on day of *C. difficile* challenge (Fig. S3A). Targeted analyses of cecal short chain fatty acid (SCFA) concentrations revealed a significant decrease in butyrate concentrations, and increase in propionate levels, in animals with IBD treated with control mAb compared to mice without IBD (Fig. S3B). Though there was a trend toward increased butyrate concentrations in animals treated with anti-p40 mAb compared to those administered control mAb, this difference was not significant (Fig. S3B). Levels of cecal bile acids were similar between all treatment groups (Fig. S3C).

The observed changes in microbiota structure in colitic mice treated with anti-p40 mAb prompted us to assess whether there is a relationship between microbiota composition and *C. difficile* susceptibility in the setting of IBD. To that end, we used Dirichlet multinomial mixtures modelling (DMM) to cluster the microbial communities in each animal into enterotypes based on the abundance of bacterial genera in their feces on the day of *C. difficile* challenge (Fig. 2D) (41). We grouped animals based on mAb treatment and subdivided them based on whether they had detectable levels of *C. difficile* in their feces one day after challenge with spores.

The DMM model with the highest likelihood (determined as having the lowest Laplace approximation value) partitioned samples into three enterotypes (Fig. 2D). Mice with communities in enterotype 1 were largely those with IBD treated with control mAb; animals with communities in enterotype 3 were solely mice in this group. Only mice treated with anti-p40 mAb had a microbiota of enterotype 2 (Fig. 2D). Examination of the top ten most abundant genera within samples revealed that, compared to enterotypes 1 and 3, enterotype 2 communities exhibited significantly higher concentrations of *Lachnospiraceae, Ruminoccoacceae,* and *Alistipes* (Fig. S4). Both enterotype 1 and 3 microbiota were characterized by increased abundances of *Enterobacteriaceae, Lactobacillus,* and *Bifidobacterium* relative to enterotype 2.

Additionally, compared to the other two enterotypes, the relative abundance of *Erysipelotrichaceae* and *Akkermansia* were increased in enterotype 1 communities (Fig. S4). Interestingly, enterotypes were significantly correlated with treatment groups and *C. difficile* susceptibility (Fig. 2D). Accordingly, all animals that carried an enterotype 3 microbiota, and a majority that harbored communities of enterotype 1, were susceptible to *C. difficile* on day 1 post-challenge (Fig. 2D). All mice with enterotype 2 communities were resistant to *C. difficile* colonization (Fig. 2D). These results indicate there is an association between microbiota structure and *C. difficile* susceptibility in the setting of intestinal inflammation.

### The microbiota from animals with active colitis is sufficient to transfer susceptibility to *C. difficile*

Our data indicate that alterations in the composition of the indigenous gut microbiota are associated with increased susceptibility to *C. difficile*. These changes largely correspond with treatment status, and thus the degree of inflammation at the time of challenge with *C. difficile* spores. To separate the development of inflammation and the associated changes in the community structure of the microbiota, we determined if the specific microbiota composition seen in colitic animals conferred susceptibility to *C. difficile* in the absence of inflammation. To do this, we conducted microbiota transfer (MT) experiments in wild-type C57BL/6 germ-free mice. Donor animals included colitic mice treated with control mAb or anti-p40 mAb. We selected donors from each treatment group that were either susceptible or resistant to *C. difficile* (Fig. 3A) to capture potential variability or features in microbiota structure that may underlie resistant and susceptible phenotypes, regardless of which treatment the animal that harbored the microbiota had received.

**Figure 3.**
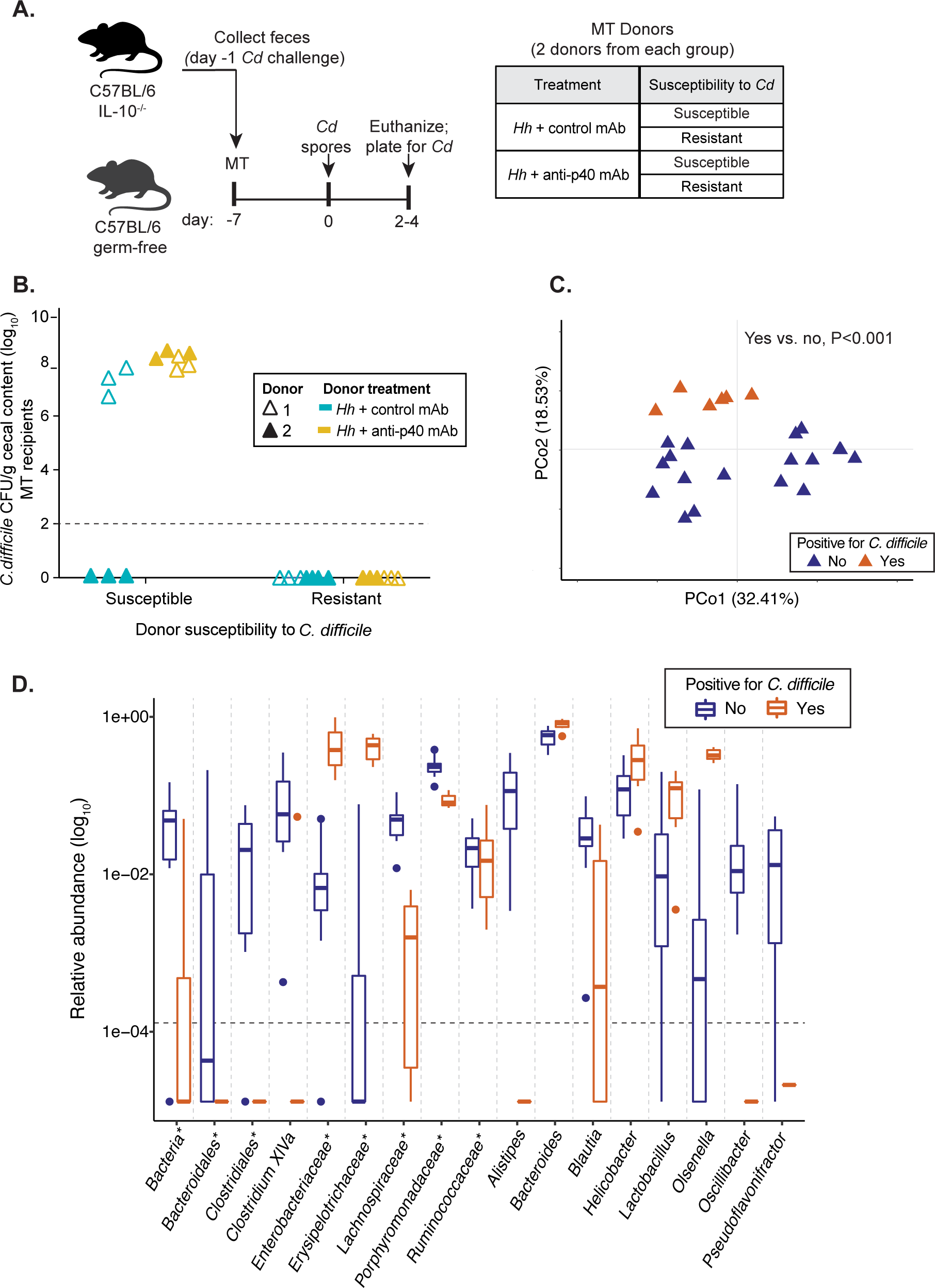
The microbiota from animals with active colitis is sufficient to transfer susceptibility to *C. difficile.* (**A**) Experimental design. Wild-type C57BL/6 germ-free mice were administered feces collected from colitic IL-10^-/-^ mice treated with control mAb or anti-p40 mAb and suspended in PBS via oral gavage. Donor feces were collected one day prior to *C. difficile* challenge; donors included animals that went on to become susceptible or remain resistant to the pathogen. Two independent experiments were completed with different donors each time (i.e., donor 1 and donor 2). One-week post-microbiota transfer (MT), recipients were challenged with *C. difficile* spores. Animals were euthanized on days 2-4 post-challenge. (**B**) *C. difficile* concentrations in the cecal contents of gnotobiotic mice at the time of sacrifice (days 2-4 post-challenge). Data are from 2 independent experiments. (**C**) Principal component analysis plot of Theta YC distances of bacterial communities in feces collected from MT recipients on day 0 of *C. difficile* challenge (7 days post-transfer). *C. difficile* positivity indicates whether animals went on to become colonized by, or remain resistant to, the pathogen by the end of the experiment. An AMOVA revealed a significant difference in microbiota structure between susceptible and resistant mice (**P<0.001). (**D**) Differentially abundant bacterial taxa in the feces of MT recipient mice on day 0 *C. difficile* challenge. Linear discriminant analysis effect size analysis (LEfSe) analysis was used to identify OTUs that were differentially abundant between recipients that were susceptible or resistant to *C. difficile*. Box plots represent mean aggregated log_10_ relative abundance of OTUs with an LDA score of ≥ 2 for each bacterial genus. Asterisks (*) denote unclassified genera. The dashed line represents the limit of detection, defined as the smallest relative abundance value in the dataset divided by 10. See also Figure S3.

Feces were collected from donor animals one day prior to *C. difficile* spore challenge (Fig. 3A) and stored until the results of challenge were known (Fig. 2D). Subsequently, the fecal pellets were thawed, suspended in PBS, and administered to recipient germ-free animals via oral gavage (Fig. 3A). The donor microbiota was allowed to engraft in recipient animals for 7 days (Fig. 3A). We used wild-type animals (IL-10^+/+^) as recipients because they do not develop colitis when colonized with *H. hepaticus* (10). This was confirmed by the low levels of lipocalin-2 detected in the stool of the MT recipients one-week post-transfer (Fig. S5A), compared to *H. hepaticus*- colonized IL-10^-/-^ SPF mice (Fig. 1B). One-week post-MT, animals were challenged with *C. difficile* spores and colonization status was determined on days 2-4 post-infection (Fig. 3A), depending on the onset of severe clinical disease (Fig. S5B).

Interestingly, recipient mice receiving MT from 3 of the 4 susceptible donors also exhibited high *C. difficile* burdens in their intestinal contents (Fig. 3B). Notably, animals positive for *C. difficile* exhibited overt disease, and the clinical scores were higher on average than those observed in IL-10^-/-^ mice with IBD and susceptible to colonization by *C. difficile* (Fig. S2A and Fig. S5C), perhaps pointing to a protective immune phenotype in animals with preexisting intestinal inflammation. Animals that received microbiota from all 4 resistant donors were subsequently resistant to *C. difficile* colonization (Fig. 3B).

Examination of the microbiota on the day of *C. difficile* challenge revealed a clear difference in community composition between susceptible and resistant MT recipients, regardless of donor susceptibility (Fig. 3C). Linear discriminant effect size (LEfSe) analysis (46) was used to identify bacterial taxa that differed significantly between MT recipients that were susceptible to *C. difficile* colonization and those that were resistant. Taxa enriched in susceptible animals included members of the *Lactobacillus, Enterobacteriaceae, Olsenella, Helicobacter, and Erysipelotrichaceae* genera; resistant animals had higher abundances of *Lachnospiraceae, Porphyromonodaceae, Clostridiales, Clostridium XIVa, Ruminoccocaceae, Alistipes,* and *Blautia,* among others (Fig. 3D). Together, these data demonstrate that the microbiota associated with IBD drives susceptibility to *C. difficile* colonization in the absence of active intestinal inflammation.

### Machine learning models predict susceptibility to *C. difficile* based on microbiota composition

Our results consistently demonstrate the critical role of the gut microbiota in regulating *C. difficile* colonization in the context of preexisting intestinal inflammation. Across experiments, we observed several taxa whose abundances consistently differed between animals susceptible or resistant to *C. difficile* colonization (e.g., *Lachnospiraceae* and *Enterobacteriaceae,* respectively). Based on these observations, we sought to determine whether there were global features of the microbiota important for conferring susceptibility to *C. difficile.* To that end, we developed random forest models to predict animals’ susceptibility to *C. difficile* colonization based on fecal microbiota composition on the day of spore challenge.

Our analysis combined samples from five independent experiments, three of which were conducted in IL-10^-/-^mice treated with control mAb or anti-p40 mAb and challenged with *C. difficile* (Figs. 1-2). The remaining two experiments involved germ-free MT mice, as outlined above (Fig. 3). In total, model development utilized 62 animals, 21 of which were susceptible to *C. difficile* colonization (34%) and 41 that were resistant (66%). To account for variation in experiment endpoints, we developed models that could classify mice as having detectable *C. difficile* in their feces one day post-challenge. Communities characterized by 16S rRNA amplicon sequence analysis were grouped by experiment and randomly partitioned into training and test datasets (65% and 35% of samples, respectively). Model performance was evaluated by measuring the area under the receiver operator characteristic curve (AUROC) (Fig. 4A-B), as well as area under the precision recall curve (AUPRC) for the test data (Fig. 4A, C). Both metrics supported high predictive performance (Fig. 4A-C).

**Figure 4.**
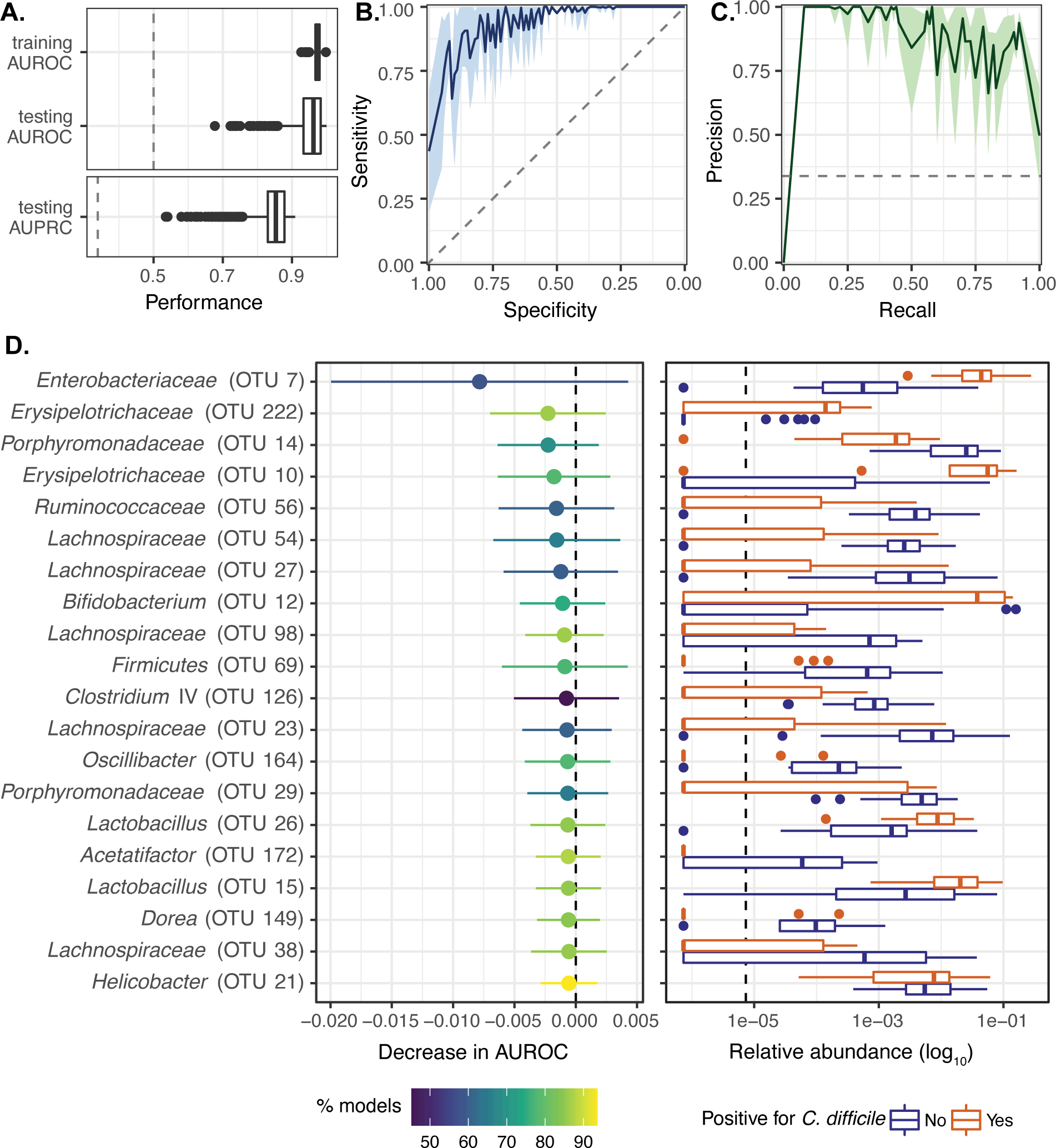
Machine learning models predict susceptibility to *C. difficile* based on microbiota composition. (**A**) Mean area under the receiver-operator characteristic curve (AUROC) on the cross-validation folds during model training, mean AUROC on the held-out test data, and mean area under the precision-recall curve (AUPRC) on the held-out test data. The dashed lines represent the baseline AUROC (0.5) and AUPRC (0.34). (**B**) Receiver-operator characteristic curve for the test data, with mean specificity plotted against sensitivity. The light green shaded area shows the standard deviation. (**C**) Precision-recall curve for the test data, with mean precision plotted against recall. The light blue shaded area shows the standard deviation. (**D**) Top 20 most important OTUs as determined by permutation feature importance (left panel). OTUs with a greater decrease in AUROC when permuted are more important. The points represent the median decrease in AUROC with the tails as the standard deviation. Color represents the percentage of models for which an OTU’s permutation AUROC was significantly different from the actual AUROC (p < 0.05). The right panel depicts the log_10_-transformed relative abundance for the top 20 most important OTUs on day 0 of the experiment, colored by *C. difficile* presence on day 1. The dashed line represents the limit of detection, defined as the smallest relative abundance value in the dataset divided by 10. See also Figure S5.

Given the good predictive value of the models, we next sought to identify operational taxonomic units (OTUs) that were most important in predicting *C. difficile* susceptibility using permutation importance (44, 47). The top 20 OTUs are depicted in Fig. 4D. OTU 7, a member of the *Enterobacteriaceae* genus, had the strongest effect on AUROC in this permutation analysis and was significant (P<0.05) in 58% of the trained models (Fig. 4D, Table S1). OTUs that significantly decreased performance for >80% of all trained models included those belonging to the *Erysipelotrichacea*e, *Lachnospiraceae*, *Lactobacillus*, *Acetatifactor, Dorea*, and *Helicobacter* genera (Fig. 4D, Table S1). Several OTUs were significant across ∼60-80% of the trained models, including those within the *Porphyromonodaceae*, *Lachnospiraceae*, *Ruminococcaeae,* and *Bifidobacterium* genera (Fig. 4D, Table S1).

Plotting the relative abundance of these top 20 OTUs in fecal samples collected from animals on the day of *C. difficile* challenge revealed clear differences between those that became positive, or remained negative, for the pathogen one day post-challenge (Fig. 4D). For example, all OTUs belonging to the *Lachnospiraceae* and *Porphyromonodaceae* genera were enriched in resistant animals whereas *Enterobacteriaceae, Lactobacillus* and *Erysipelotrichaceae* were more abundant in susceptible mice. These findings highlight specific microbiota taxa as important for modulating susceptibility to *C. difficile* colonization.

## Discussion

The indigenous gut microbiota plays a central role in colonization resistance to *C. difficile,* in part through interactions with the host (8–11, 45). Murine models have been critical for advancing our understanding of the structural and functional aspects of the microbiota that regulate susceptibility to CDI (48). However, most murine studies have focused on *C. difficile* colonization in the context of antibiotic treatment (4, 24). There is a need to develop appropriate models to understand mechanisms underlying non-antibiotic risk factors for CDI development, including underlying IBD.

Here, we demonstrate that intestinal inflammation alters microbial community structure in mice with colitis, and that these changes are associated with a loss of colonization resistance to *C. difficile.* We show that inflammation sculpts the microbiota to permit *C. difficile* colonization, though it is not required in addition to, or instead of, these alterations. These findings suggest a temporal relationship between the development of inflammation and loss of microbiota-mediated resistance to *C. difficile.* Collectively, our results provide a new perspective on the relative contributions of the host and microbiota in colonization resistance to *C. difficile*. Importantly, they support the primacy of the microbiota in regulating *C. difficile* susceptibility, as described for antibiotic-associated CDI pathogenesis.

We provide a novel system to study features of the microbial community modulating vulnerability to CDI in the setting of intestinal inflammation. Notably, many of the taxa we consistently identified as associating with susceptibility to *C. difficile* colonization have previously been implicated in CDI. For example, in agreement with our studies, higher concentrations of *Lactobacillaceae, Erysipelotrichaceae,* and *Enterobacteriaceae* are generally associated with susceptibility to CDI in murine and human systems (3, 35, 49, 50). In contrast, *Lachnospiraceae, Ruminococcaceae,* and *Porphyromonodaceae,* which were enriched in resistant animals, are linked with protection against, or clearance of, *C. difficile* in mouse models of CDI (3, 50–54).

Regarding infection in humans, these taxa are more abundant in healthy people compared to patients with CDI (49). Our findings, in conjunction with these studies, suggest there are conserved microbial signatures that correlate with susceptibility to CDI in various intestinal contexts. Delineating the mechanisms by which these key taxa, and the microbial community as a whole, modulate *C. difficile* colonization will be an important avenue for future investigations.

To that end, several taxa associated with resistance to CDI in our study and others (e.g., *Lachnospiraceae* and *Ruminococcaceae)*shape the intestinal metabolic milieu in ways that could interfere with *C. difficile* colonization, such as through the production of SCFAs and secondary bile acids (3, 55–60). Previous work has demonstrated that antibiotics alter intestinal concentrations of these compounds (59, 61, 62) and these alterations, particularly in bile acid metabolism, support *C. difficile* growth (59). Interestingly, we found no significant differences in cecal SCFA and bile acid profiles between colitic mice treated with control mAb and anti-p40 mAb, despite variations in microbiota structure and *C. difficile* susceptibility observed between these groups. These data suggest that, in this system, susceptibility to CDI is likely influenced by factors other than, or in addition to, these compounds. Thus, it is becoming clear that the relationship between *C. difficile* the intestinal environment, as it is shaped by both the microbiota and the host, is a complex, multilayered system of multiple interactions.

From a clinical standpoint, our findings are intriguing considering the intersection between IBD and risk for CDI (22). People with IBD have altered gut microbial community structure compared to healthy individuals (13). Accordingly, many of the differences in microbiota composition observed in patients with IBD, such as loss of *Lachnospiraceae* and enrichment in *Enterobacteriaceae* species, correspond with those described here, and by others, as being linked with CDI development (63, 64). Thus, our study suggests the inflammatory landscape shaped by IBD enriches for certain susceptibility-associated taxa while hindering expansion of resistance-associated bacteria. Therefore, controlling IBD-associated inflammation may alter the microbiota in ways that restrict *C. difficile* growth. Further explorations of the host-microbiota-CDI interface in the setting of IBD in murine models and patients will reveal context-dependent mechanisms of *C. difficile* pathogenesis most relevant to patients with IBD. These studies are critical in the development of prophylactic and therapeutic strategies for managing CDI in IBD patients.

## Acknowledgments

We acknowledge the University of Michigan Germ-Free Mouse Core, Microbiome Core, and Metabolomics Core for assistance with data curation, as well as McClinchey Histology Labs, Inc. (http://www.mhistolab.com) for preparing histology slides. We also thank Helen Warheit-Niemi and other members of Dr. Bethany Moore’s laboratory for sharing equipment as well as providing primers and analysis templates for RT-PCR experiments. Lastly, thank you to Dr. Christine Bassis for guidance and input with troubleshooting 16S rRNA amplicon sequencing analyses. This work was supported by the National Institutes of Health cooperative agreement AI124255 to V.B.Y. and P.D.S. and AI007528 to M.R.B. L.A.C was supported by grant number UL1TR002240 from the National Center for Advancing Translational Sciences (NCATS).

## Author contributions

M.R.B conceptualized the study, curated and analyzed the data, and wrote the manuscript. K.L.S. conducted the machine learning analyses and helped interpret the results, as well as assisted with writing the Methods section. L.A.C. advised on project conceptualization and, along with K.C.V and A.K.S., assisted with mouse experiments. I.L.B. conducted histopathological analyses. P.D.S. advised on DMM and machine learning analyses while V.B.Y. helped with study conceptualization, supervised the work, and revised and edited the manuscript. All authors critically revised the manuscript and approved the final version to be published.

## Supplemental material titles and legends

**Table S1 Primers and probes used for PCR and RT-PCR analyses**

**Figure S1 Histopathological and gene expression analyses of colon tissue from IL-10^-/-^ mice without IBD and mice with IBD treated with control mAb or anti-p40 mAb.** (A) Experimental timeline. Animals were inoculated with *H. hepaticus (Hh)* or sterile tryptic soy broth (non-IBD controls) via oral gavage. Fourteen days later (day 0), anti-p40 mAb, isotype control mAb, or vehicle were administered to mice via intraperitoneal (IP) injection every 3-4 days for 3 weeks. Mice were then euthanized and colon tissue collected for histological and RT-PCR analyses. (**B)** Histopathological damage in the colons collected from mice without colitis and with colitis treated with control or anti-p40 mAb. The degree of lymphocytic inflammation was determined using a 4-point scale. Data from 2 independent experiments were analyzed using a Kruskal Wallis test followed by the Dunn’s multiple comparison’s test (** P<0.01, **** P<0.0001). **(C)** Colon tissue collected from mice colonized with *H. hepaticus* or mock colonized with sterile broth after 3 weeks of antibody treatment. Representative hematoxylin and eosin images are shown. (**D-G)** RT-PCR analysis of genes downstream of p40-mediated signaling pathways in colon tissue collected from mice with and without IBD after 3 weeks of treatment with mAb or vehicle. Results are from 2 independent experiments. Expression of β-actin was used to normalize RNA in samples. Fold change was calculated relative to a sample within the “*Hh* + control mAb” group for each gene. Statistical significance was determined via an ANOVA followed by Tukey’s test (D, E, F) or Kruskal Wallis with Dunn’s multiple comparison’s test (G) (*P<0.05, **P<0.01, ***P<0.001, ****P<0.0001).

**Figure S2 Clinical and histopathological scores of mice with IBD and treated with control mAb or anti-p40 mAb and challenged with *C. difficile.*** (A) Clinical disease of mice with colitis on day 9 post-challenge with *C. difficile* (day of sacrifice; animals are the same as those depicted in Fig. 1 in the main text). Disease severity did not significantly differ between animals that were positive or negative for *C. difficile* at any point throughout the experiments within each treatment group, or between groups. A Kruskal Wallis test followed by the Dunn’s multiple comparison’s test was performed. (**B**) Cecal and colon histopathology scores (composite score of edema, epithelial damage, and inflammatory cell infiltration) in mice with IBD treated with control mAb or anti-p40 mAb and challenged with *C. difficile*. *Hh = H. hepaticus, Cd = C. difficile*.

**Figure S3 SCFA and bile acid concentrations in cecal contents of mice without IBD and mice with IBD treated with control or anti-p40 mAb.** (A) Principal component analysis plot of Theta YC distances of bacterial communities in cecal contents collected from mice without colitis and with colitis after 3 weeks of receiving control mAb or anti-p40 mAb. (B) Concentrations of acetate, butyrate, and propionate in cecal contents of mice without IBD and with IBD treated with control mAb or anti-p40 mAb, as measured via LC-MS. Results for each compound were analyzed using an ANOVA followed by Tukey’s test (*P<0.05, **P<0.01). (C) LC-MS analysis of primary and secondary bile acids in cecal contents. Statistical significance was determined via an ANOVA followed by Tukey’s test or Kruskal Wallis with Dunn’s multiple comparison’s test depending on data distribution (*P<0.05). Asterisked bile acids denote those only produced in mice. CA = Cholic acid; αMCA= Alpha muricholic acid; βMCA= Beta muricholic acid; GCA = Glychocholic acid; TCA=Taurocholic acid, UDCA = Ursodeoxycholic acid; DCA = Deoxycholic acid; HDCA= Hyodeoxycholic acid, *w*MCA = Omega muricholic acid.

**Figure S4 Abundance of top ten bacterial taxa in feces of mice with gut microbial communities of enterotypes 1, 2, and 3.** Log_10_ relative abundance of top ten most abundant taxa in fecal samples collected from animals on the day of *C. difficile* challenge and colored by enterotype (Fig. 2D in main text). Box plots represent mean aggregated relative abundance of OTUs within each bacterial genus. Asterisks (*) denote unclassified genera. Dashed line represents the limit of detection, defined as the smallest relative abundance value in the dataset divided by 10. Data were analyzed via a Kruskal Wallis and Dunn’s test for each genus (*P<0.05, **P<0.01, ***P<0.001, ****P<0.0001).

**Figure S5 Fecal lipocalin-2 concentrations, survival, and clinical scores of microbiota transfer (MT) recipients.** (**A**) Fecal lipocalin-2 levels of MT recipient mice at baseline and 7 days post-transplant. Mice exhibited lower lipocalin-2 concentrations relative to IL-10^-/-^ mice with IBD (Fig. 1B in main text). (**B**) Survival curve of MT recipients. Several animals harboring microbiota from *C. difficile* susceptible donors died prior to the experiment endpoint. (**C**) Clinical disease of MT recipients at the time of sacrifice. Animals colonized by *C. difficile* exhibited overt signs of disease. *Hh = H. hepaticus, Cd = C. difficile*.

Table S2 Feature Importance analyses, top 20 OTUs (corresponds with Fig. 4D in main text). OTUs are ranked by mean decrease in AUROC after permutation.

